# A High Performance, Inexpensive Setup for Simultaneous Multisite Recording of Electrophysiological Signals and Wide-Field Optical Imaging in the Mouse Cortex

**DOI:** 10.1101/177188

**Authors:** Edgar Bermudez Contreras, Sergey Chekhov, Jennifer Tarnowsky, Jianjun Sun, Bruce L. McNaughton, Majid H. Mohajerani

## Abstract

Simultaneous recording of optical and electrophysiological signals from multiple cortical areas may provide crucial information to expand our understanding of cortical function. However, the insertion of multiple electrodes into the brain may compromise optical imaging by both restricting the field of view, and interfering with the approaches used to stabilize the specimen. Existing methods that combine electrophysiological recording and optical imaging *in vivo* implement either multiple surface electrodes or a single electrode for deeper recordings. To address such limitation, we built a microelectrode array (hyperdrive) compatible with wide-field imaging that allows insertion of up to 12 probes into a large brain area (8 mm diameter). The developed hyperdrive is comprised of a circle of individual microdrives where probes are positioned at an angle leaving a large brain area unobstructed for wide-field imaging. Multiple tetrodes and voltage-sensitive dye imaging (VSDI) were used for simultaneous registration of spontaneous and evoked cortical activity. The electrophysiological signals were used to extract local field potential (LFP) traces, multiunit and single-unit spiking activity. To demonstrate our approach, we compared LFP and VSD signals over multiple regions of the cortex and analyzed the relationship between single-unit and global cortical population activities. The study of the interactions between cortical activity at local and global scales, such as the one presented in this work, can help to expand our knowledge of brain function.

## 1. Introduction

A fundamental goal of neuroscience is to understand the underlying mechanisms that are employed by the brain to process information. However, in a complex system such as the brain, it is difficult to explain the behaviour of the system by only studying its components in isolation. Rather, it is crucial to understand how the interactions of the components give rise to the behaviour of the system. Analogously, in order to understand the brain dynamics, it is necessary to analyze its activity at different scales. As we know, the behaviour of a neuronal network is not only determined by its connectivity but also by the external inputs which might involve multiple and distant networks. Therefore, in order to fully understand how the brain process information, it is necessary to be able to study neuronal activity at local and global spatiotemporal scales (Luczak et al. 2015; McVea et al. 2016; Sofroniew et al. 2016; Chen and Wilson 2017; Thome et al. 2017; Xiao et al. 2017; Zhuang et al. 2017).

Electrophysiological recordings have been used extensively to study neuronal activity, and with the development of tetrode arrangements, it has become an invaluable tool to monitor spiking activity of individual cells at any brain depth (McNaughton et al. 1983). The overall reliability of the technique (Schmidt et al. 1976; Kralik et al. 2001; Moxon et al. 2009; Schwindel et al. 2014) provides easy transfer of acquired experimental data into scientific knowledge. However, technical problems that occur when implanting highly dense electrode arrays or theoretical difficulties in determining the signal sources make this technique difficult to apply for recordings over large areas of the cortex (Battaglia et al. 2004; Kajikawa and Schroeder 2011; Buzsáki et al. 2012).

Optical methods, however, offer both sufficient temporal and spatial resolution for real-time analysis of brain processing (Grinvald et al. 2001; Frostig 2009). Wide-field optical imaging and, in particular, voltage-sensitive dye imaging technology has evolved into a convenient tool to study neuronal activity dynamics over large areas of the cortex with high temporal and spatial resolution (Petersen et al. 2003; Grinvald and Hildesheim 2004; Mohajerani et al. 2011, 2013; Geng and Wu 2017). The spatial resolution reaches 25-65 μm per pixel while the size of imaged brain area is sufficient to record electrical activity of most of the mouse cortex (Lim et al, 2015), 1 cm^2^ area monkey’s cortex (Yang & Heeger, 2007) or 0.3 cm^2^ area over the cat cortex (Benucci et al, 2007). With these characteristics, VSDI is a great technique to study neuronal dynamics over large cortical areas. However, VSDI has limitations that are important for a complete study of the complex interactions of neuronal networks. In particular, VSDI mainly captures subthreshold neuronal activity located within different cortical layers (Grinvald and Hildesheim 2004; Mohajerani et al. 2010; Chemla et al. 2017). Therefore, wide-field optical imaging and VSDI in particular, are great candidates to be combined with multi-site electrophysiological recordings. However, other recently developed longitudinal mesoscale imaging options such as Calcium (GCaMP) or glutamate sensors (iGluSnFR) which reflect neuronal activity at the population level can also be employed for similar purposes (Xie et al. 2016).

The idea of combining electrophysiological recordings with VSDI is not new. There are already approaches available to image with parallel cell recordings from the acute brain slices (Tominaga et al. 2001; Carlson and Coulter 2008; Ayzenshtat et al. 2010; Mapelli et al. 2010; Ferrea et al. 2012). However, such studies are limited to slice preparations. Additionally, simultaneous single-unit recordings and VSDI *in vivo* have been pioneered by Grinvald’s group (Grinvald et al. 2001; Grinvald and Hildesheim 2004) but their work is limited to a single region in the brain. As well, there have been approaches combining VSDI over large cortical areas and electrophysiological recordings (Ferezou et al. 2007; Ayzenshtat et al. 2010; Mohajerani et al. 2013) but, in these cases, they employ a small number of surface or pipette electrodes that are not suitable for recording signals from deep brain structures or from multiple units. The attempts to combine VSDI with deep recordings *in vivo* include experiments on a very well-studied model, primate V1, with separate preparations for each method (Chen et al, 2012), as well as whole-cell recordings with simultaneous VSDI to describe the propagation of excitation in the rat barrel cortex (Petersen et al. 2003; Berger et al. 2007). However, an approach to combine large scale VSD imaging with multisite deep recordings of extracellular electric potentials in rodent cortex remained to be developed. Recently, (Kunori and Takashima 2015) developed a transparent multielectrode array that registers field potentials from the brain surface *in vivo* that can be combined with brain imaging. However, their approach appears unsuitable for deep multi-unit recordings.

In this article, we present a novel methodology to simultaneously monitor neuronal activity at two different spatial scales and of a different nature. We record local cortical activity over multiple areas using multisite electrodes and global cortical activity using wide-field VSD imaging. To demonstrate our approach, we compare cortical activity simultaneously recorded by VSD imaging with local field potentials (LFP) and single-unit spiking activity (SUA) and study their relationship. The study of how these signals interact, as facilitated by our hyperdrive, can potentially expand our understanding of information processing at micro-and mesoscales, which in turn is crucial to study brain function.

## 2. Materials and Methods

### Animals

All experiments were carried out on adult (20-30 g, age 2-4month) wild type C57/Bl6 mice (n = 5) or B6.Cg-Tg (Thy1-COP4/EYFP) 18Gfng/J mice (n = 2). Mice were housed under standard conditions, in clear plastic cages under 12 h light, 12 h dark cycles. Mice were given ad libitum access to water and standard laboratory mouse diet at all times. All protocols were approved by the Animal Welfare Committee of the University of Lethbridge and were in accordance with guidelines set forth by the Canadian Council for Animal Care.

### Surgery

Mice were anesthetized with 15% urethane (1250 mg/kg) and fixed in a stereotactic apparatus. Body temperature was maintained at 37°C with an electric heating pad regulated by a feedback thermistor. Mice were given Dexamethasone (80 μg) intramuscularly to prevent inflammation and Lidocaine (50 μl, at 2%) into the area of the skin incision over the skull. The plastic head plate (inner diameter 8 mm) was attached to the bone with dental cement (Osborne and Dudman 2014). A ∼8mm diameter single cranial window was made over both mouse cortical hemispheres (2.5-5.5 mm, anterior-posterior and 0–4 mm laterally from bregma) using a high-speed dental drill (Kyweriga et al. 2017). To keep the brain cool, the drilling was done intermittently and the skull was moistened with brain buffer. Caution was given to keep the dura intact when removing the bone. For each hour under anesthesia, the mouse was given 10 ml/kg of 20 nM glucose in brain buffer IP to maintain hydration. A tracheotomy was performed for better ventilation.

The setups to fixate an animal and a hyperdrive were similar to RIVETS, described in (Osborne and Dudman 2014). When the animal’s head was secured between the plastic forks, the hyperdrive was centered above the head plate. Tetrodes were carefully loaded down below the brain surface and traveled about 600-750 μm at an angle of about 45°. The tetrodes easily penetrated the brain surface when *dura mater* was removed (Figure 2B), but also were able to pass through the intact *dura mater.* Finally, one of the tetrodes was placed above the cortex surface to serve as a reference. The mouse was then placed on a metal plate that could be mounted on the stage of the upright macroscope, and the skull was secured using RIVETS fork system. A modified fork system was designed to hold the electrode array. Then the animal with the recording setup was then transferred to the experimental table and placed under the VSDI camera (Figure 2A).

### Electrophysiological recordings

We used a custom 3D-printed plastic hyperdrive (similar to the electrode array first described in Gothard, 1996) consisting of 12 slots for individually movable microdrive probes (Figure 1). Each microdrive can be loaded with a tetrode, stimulating electrode, sharp metal electrode with glass/plastic coating (Reitboeck 1983; Steenland and Mcnaughton 2015) or fiber optic. The hyperdrive implemented several unique features to allow for simultaneous wide-field optical imaging. The features are the following: a 7.5 mm circular opening in the center provides sufficient brain area to image (Figures 1A and 2B); a miniature plastic edge was designed to mount an 8 mm cover glass (Figure 1B). The cover glass stays above the tetrodes, protecting the brain surface and reducing brain pulsations (Helmchen and Konnerth 2011); the slots for microdrives were made slightly curved to reduce the overall height of the hyperdrive and facilitate illumination of the brain surface for optical imaging (Figure 1B); metal hexnut glued into the plastic to provide higher precision and longevity of the hyperdrive (Figure 1B).

**Figure 1.**
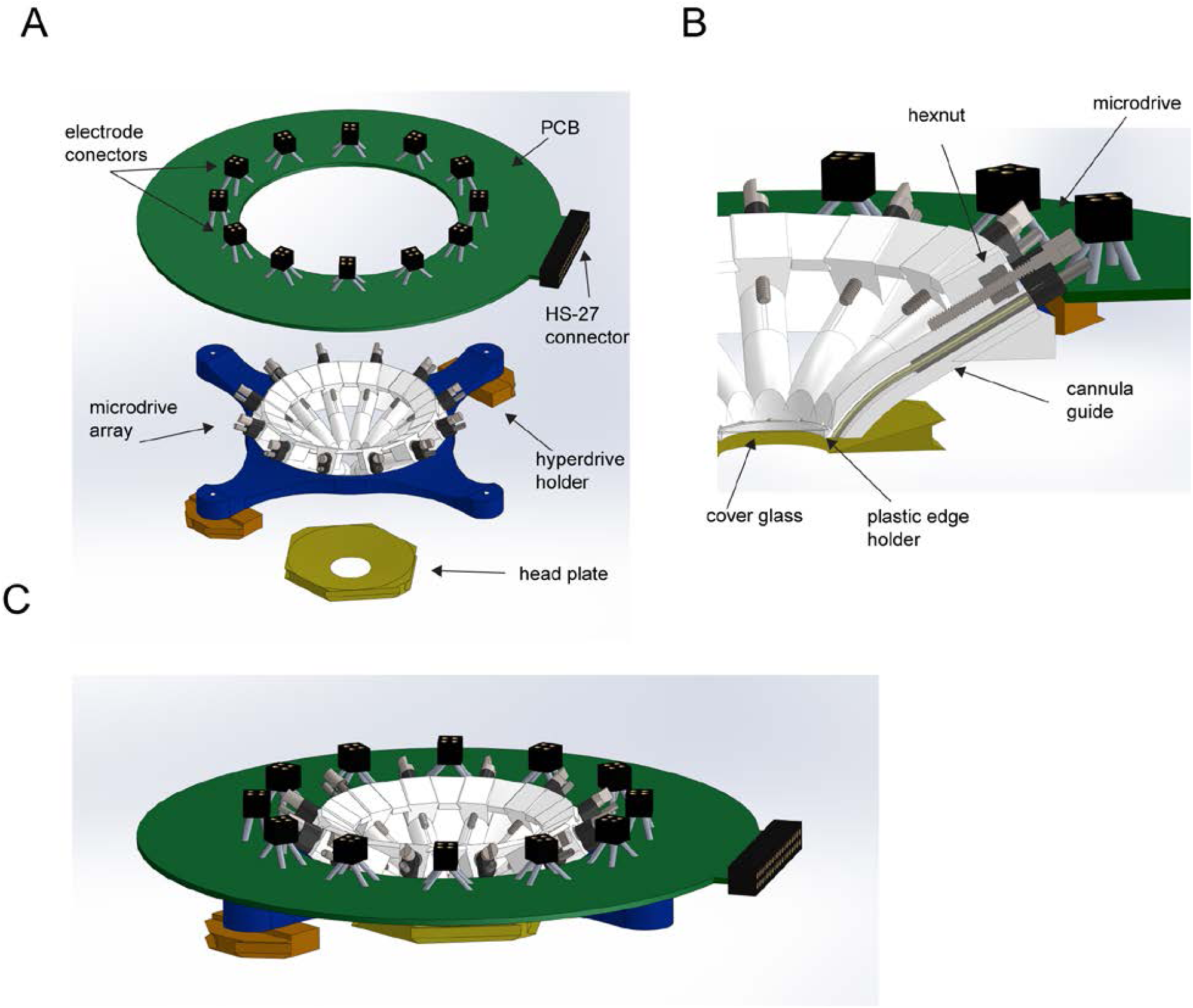
Overview of the wide-field imaging hyperdrive. (A) Renderings of the main parts of the hyperdrive: Printed circuit board (PCB); electrode connectors; HS-27 connector (Neuralinx); circular microdrive array; hyperdrive holders; headplate. (B) Cross section of the microdrive. The metal screw is moving inside the hexnuts glued into the plastic base. The tetrode (not shown) is glued to the microdrive and moves inside the guide cannula, which opens below the coverslip glass. (C) Rendering of the assembled hyperdrive.

The tetrodes that were used comprised of 4 twisted 12.5 μm nichrome wires with polymide coating (Sandvik, USA) and were gold plated to reduce the impedance to 100 kΩ or lower. The individual tetrode wires were soldered into a custom designed ‘Flex-connector’ (NeuroTek) that attached to the PCB with a Mill-max connector (Figure 2C). A custom built circuit board on the top of the hyperdrive was connected to a unity-gain headstage (HS-27, Neuralynx, Bozeman, MT) to provide a low-noise signal transition to the recording system (Figure 1C and 2C). The signal was recorded and time stamped by Digital Lynx 16 SX system (Neuralynx, Bozeman, MT). A reference electrode was placed above the cortex so that the tip was submerged either into the brain buffer or the agarose.

**Figure 2.**
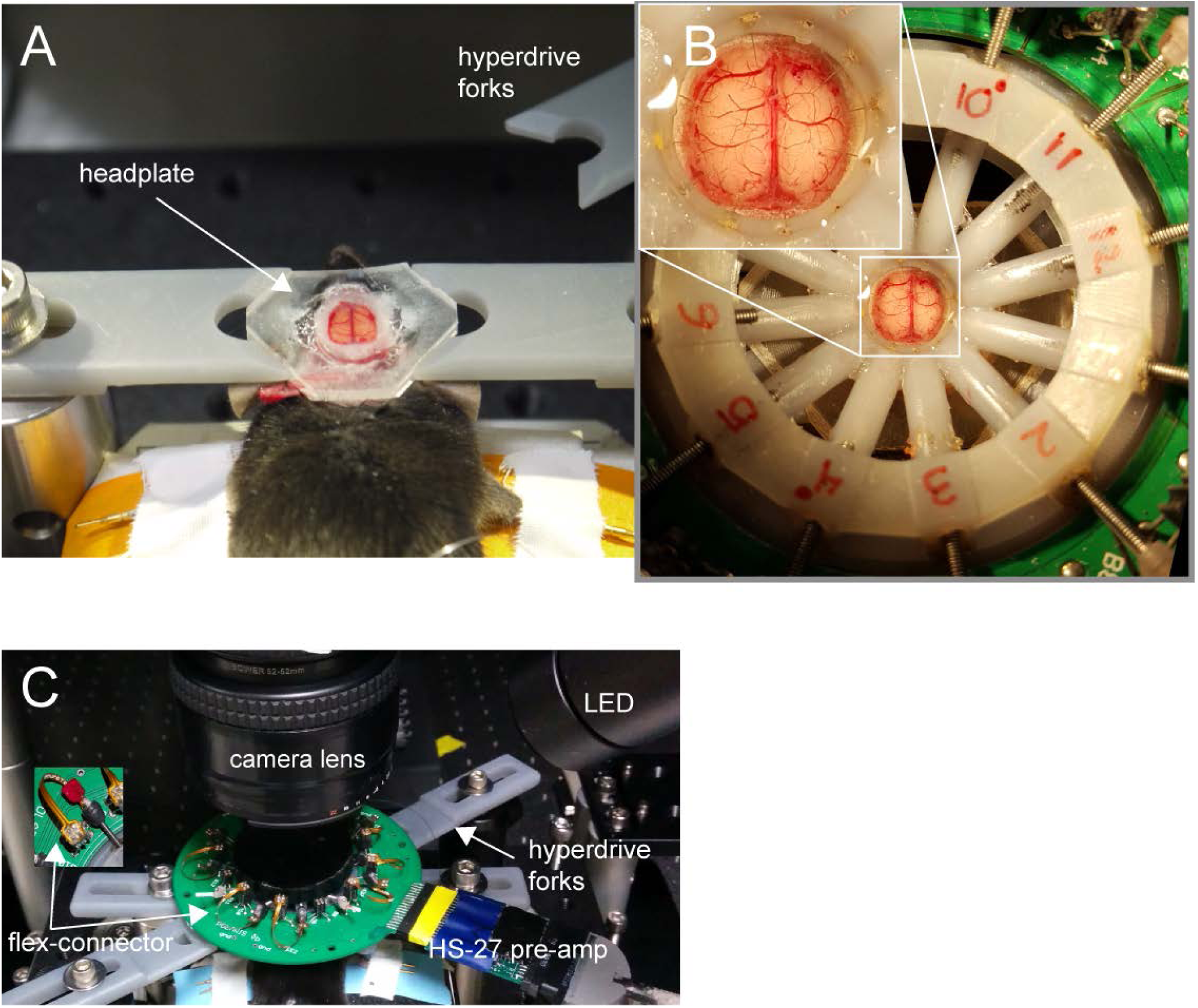
Hyperdrive and wide-field optical imaging preparation. (A) A urethane anesthetized mouse with bilateral craniotomy is fixed between two forks using glued in headplate. (B) Hyperdrive centered above bilateral craniotomy. Inset: Enlarged view of the bilateral craniotomy with the electrodes inserted into the brain. (C) Over view of the experimental setup. Inset: flex-connectors.

To record neuronal spiking activity, the extracellular electric signal was high-pass filtered (0.1 Hz), amplified 1,000 times and digitized at 32 kHz using a Digital Lynx 16 SX system and an HS-27 headstage (Neuralynx, Bozeman, MT). LFP traces were recorded from the same tetrodes and down-sampled at 312 Hz for analysis. Spike sorting was performed semi-automatically using Klustakwik (http://klustakwik.sourceforge.net), followed by manual clustering using Mclust (https://cran.r-project.org/web/packages/mclust/index.html). Histology suggests that the tetrodes tips were located in layers IV, V in the cortex (data not shown).

### VSD imaging

After inserting the tetrodes to the target sites, the dye RH-1691 (optical Imaging, New York, NY) was dissolved in brain buffer solution (0.5 mg/ml) and applied to the exposed cortex for 60-90 min. After washing unbounded dye for 5-10 min with brain buffer solution, the brain was covered with 1.5% agarose made in HEPES-buffered saline and sealed with a glass coverslip. This procedure reduced the movement artifacts produced by respiration and heartbeat. For VSD data collection, 12-bit images were captured at 150 Hz during evoked activity and at 100 Hz during spontaneous activity with a charge-coupled device (CCD) camera (1M60 Pantera, Dalsa, Waterloo, ON) and an EPIX E8 frame grabber with XCAP 3.8 imaging software (EPIX, Inc., Buffalo Grove, IL). The dye was excited using a red LED (Luxeon K2, 627 nm center) and excitation filters of 630 ± 15 nm. Images were taken through a macroscope composed of front-to-front video lenses (8.6 x 8.6 mm field of view, 67 μm per pixel). The depth of field of our imaging setup was 1 mm. Reflected VSD fluorescence was filtered using a 673-to-703 nm bandpass optical filter (Semrock, New York, NY). To reduce potential VSD signal distortion caused by the presence of large cortical blood vessels, we focused into the cortex to a depth of ∼ 1mm.

### Evoked and spontaneous activity

For sensory-evoked activity, we recorded baseline activity for 900 ms and 4100 ms after a single 1 ms electrical pulse (300 μA) was delivered to the hind left paw for each trial. Because animal brain states show spontaneous changes, we averaged 20 trials of stimulus presentation to reduce these effects. To correct for time-dependent changes in VSD signals due to bleaching artifact, we also collected 20 non-stimulation trials that were used for normalization of the evoked data. A 10-s interval between each sensory stimulation was used. In a previous work, VSD fluorescence was measured across the cortex using histology and demonstrated relatively high labeling at a depth of ∼750 μm (Mohajerani et al. 2010). Nonetheless, to reduce regional bias in VSD signal caused by uneven dye loading or brain curvature, all VSD responses were expressed as a percentage change relative to baseline VSD responses (ΔF/F_0_ × 100%) using ®Matlab (Mathworks, Natick, MA). VSD imaging of spontaneous activity was continuously recorded in the absence of sensory stimulation for 15 min period with 10 ms (100 Hz) temporal resolution. Slow, time-dependent reductions in VSD fluorescence were corrected in Matlab using a zero-phase lag Chebyshev bandpass filter (zero-phase filter) at 0.1–6 Hz. Ambient light resulting from VSD excitation (630 nm) was measured at 8.65 × 10^−3^ W/m^2^. The total duration of the VSD excitation in a typical imaging experiment ranged from 900 to 1,200 s. The fluorescence changes were quantified as (F–F_0_)/F_0_ × 100, where F is the fluorescence signal at any given time and F_0_ is the average of fluorescence over baseline frames. To analyze the relationship between single-unit activity (SUA) and neuronal population activity we calculated the spike-triggered average VSD (STA maps) of each neuron by taking the mean of the VSD signal over all the times when such neuron fired (Xiao et al. 2017).

### VSDI and electrophysiological signals synchronization and comparison

VSD images and electrophysiological records were digitized on two separate systems with different sampling rates (200 Hz and 32 kHz, respectively). To synchronize these signals we recorded the clock from the EPIX frame grabber, the excitation LED trigger and the electrical stimulation signals in the Digital Lynx 16 SX system (Neuralynx, Bozeman, MT) via the TTL port. During off-line analysis, we used such signals to align imaging and electrophysiological data.

We compared the LFP and VSD signals using the Pearson correlation coefficient during evoked and spontaneous activity. During evoked activity, we divided the signals into three periods: baseline, early and late responses. Baseline activity consisted of activity before the stimulus onset (900 ms). Early evoked response consisted of the first 250 ms after stimulus onset. Late evoked responses were considered as the following 250 ms after the early evoked response. For the spontaneous activity, we calculated the similarity between LFP and VSD signals as the correlation coefficient during 15 min of spontaneous activity divided into segments of 1 sec (used to calculate the mean similarity between the signals).

## 3. Results

### Combined VSD imaging and multi-site electrophysiological recording in response to the sensory-evoked stimulation

Using a preparation with a bilateral craniotomy that exposed a large portion of the dorsal cortex in both hemispheres (Figure 3A), we were able to monitor brain activity using both VSDI and electrophysiology simultaneously.

**Figure 3:**
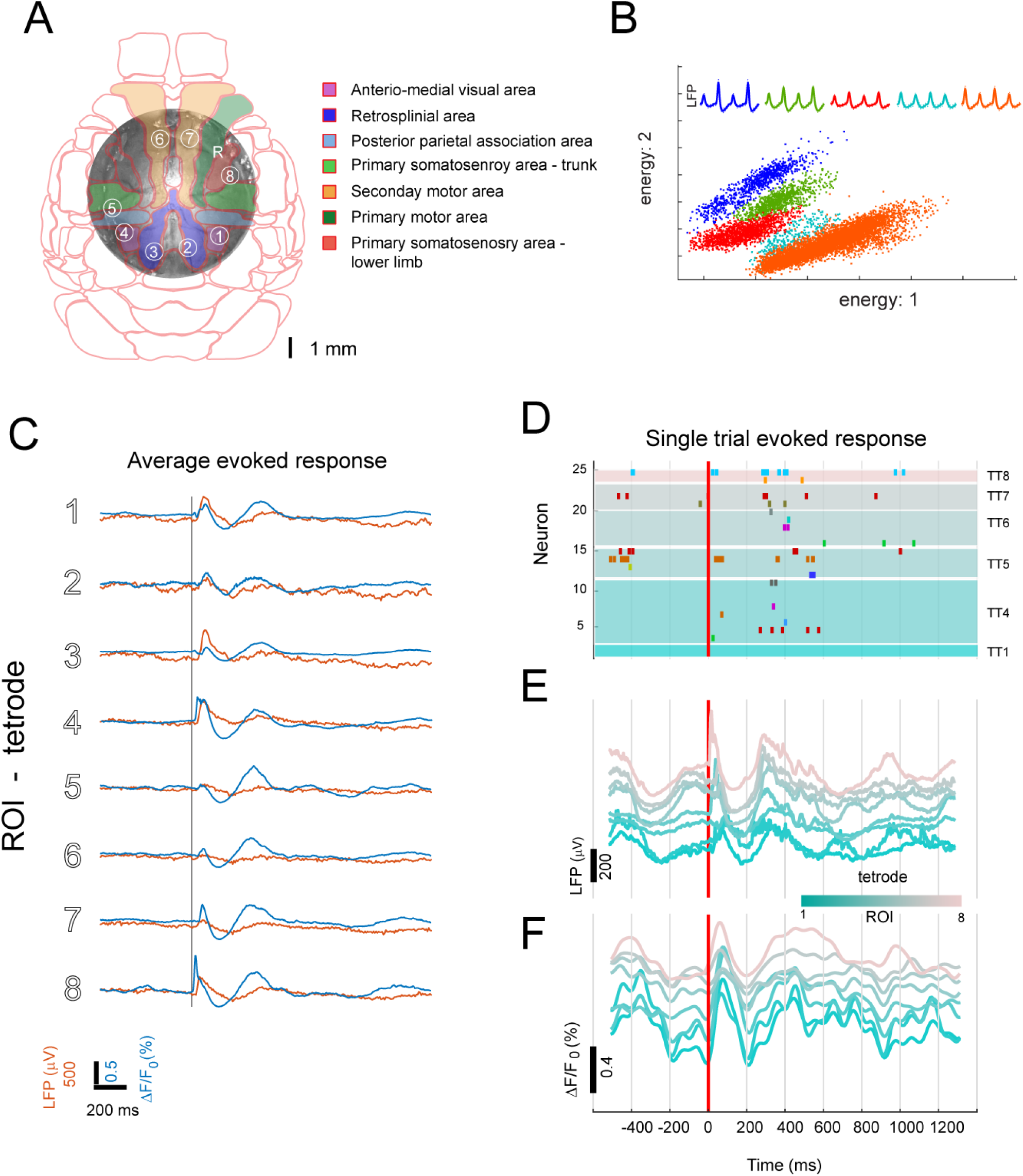
Evoked response to sensory stimulation using electrophysiology and VSDI simultaneously. (A) Photomicrograph of wide bilateral craniotomy preparation with the position of the electrodes marked as numbered circles in reference to the Allen Brain Institute Mouse Brain Atlas. (B)Example of five putative pyramidal neurons recorded in one tetrode. (C) Average evoked VSD (blue) and LFP (red) response to electrical stimulation of the hind paw. The gray vertical line represents the stimulus onset. (D) Single trial raster plot of single unit activity in response to one pulse of electrical stimulation of the left hind paw. Different colors represent different neurons (E) LFP evoked response to the same stimulation trial as in (D). Each line represents the LFP from one of the 8 electrodes. (F) VSD evoked response to the same stimulation trial as in (D). Each line represents the VSD signal from one of the 8 ROIs in (A).

To compare VSD and electrophysiological evoked responses, we averaged the VSDI signal within regions of interest (ROI) of 5 pixel diameter around the point where each tetrode was inserted into the cortex (Figure 3A). When stimulating the hind paw of lightly anesthetized mice, we observed unique patterns of cortical depolarization (Figure 4D). Consistent with previous studies (Mohajerani et al., 2011; Lim et al., 2012 and 2014), we found that brief electrical stimulation of left hindpaw led to activation of contralateral primary hindlimb (HL) somatosensory cortex around 20-30ms after stimulus onset. The activation of contralateral HL cortex was followed by an expansion of depolarization within the contralateral hemisphere into neighboring areas. In addition, an activation of primary hindlimb cortex within the ipsilateral hemisphere (Figure 4D) follows shortly after the initial contralateral response. The average temporal profiles of the evoked response in both VSDI and electrophysiology are similar for most of the ROIs (Figure 3C). However, for ROIs near HL somatosensory cortex (ROIs 1, 5 and 8), the latency of response is shorter than in other ROIs. Figure 3B shows an example of sorted putative pyramidal neurons from a single tetrode. In a single trial we can observe that evoked response to a single pulse of HL electrical stimulation expands over large areas of the cortex (Figure 3D-F).

**Figure 4.**
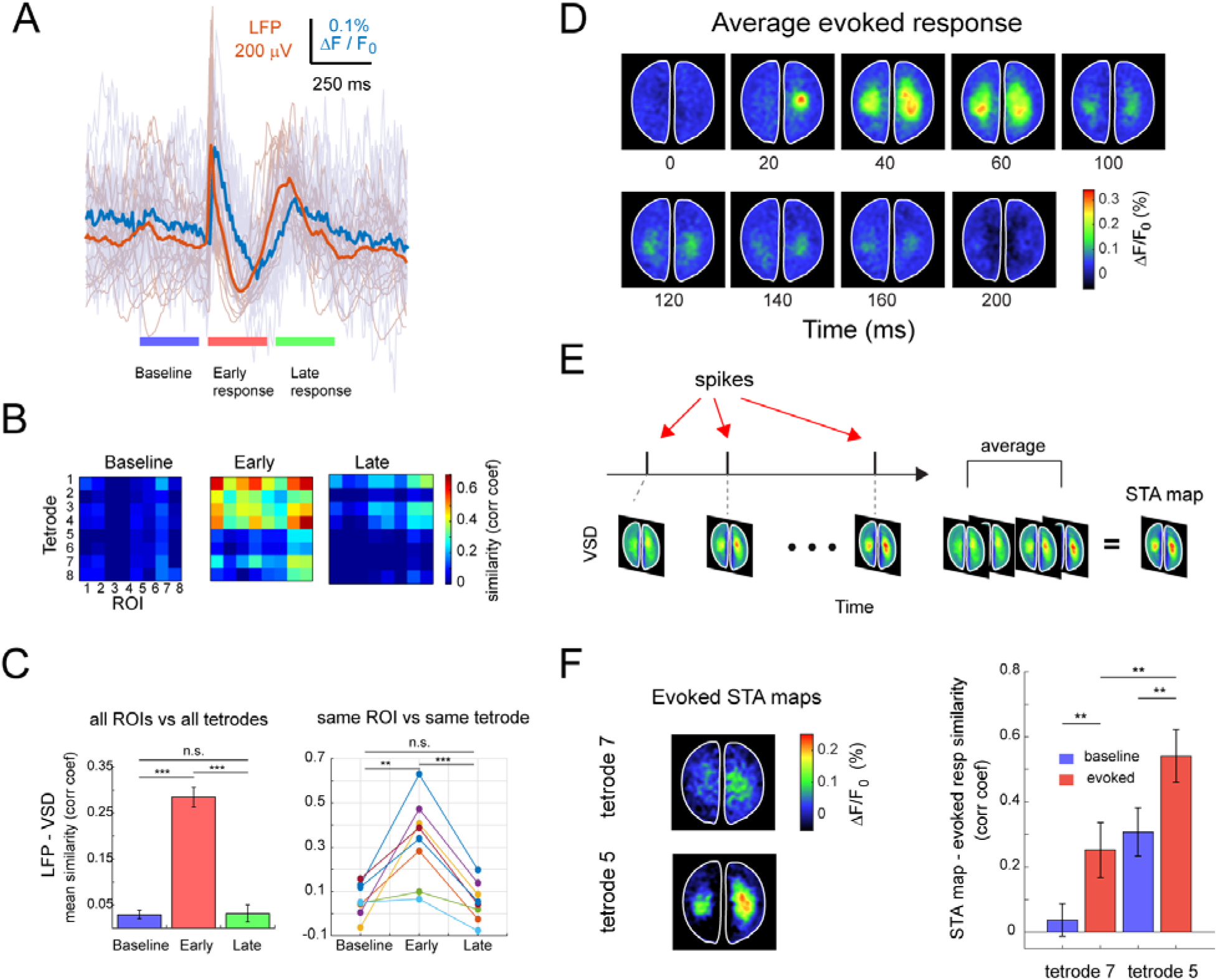
Relationship between local field potential, single unit activity and voltage sensitive dye signals during evoked responses. (A) Example of mean LFP (orange) from one tetrode and single trials (shaded) and mean VSD (blue) from same ROI and single trials (shaded). Evoked responses are divided into three periods: baseline (last 250 ms before stimulus onset), early evoked response (first 250 ms after stimulus onset) and late evoked response (250 ms after the evoked1 period). (B) Similarity matrices (correlation coefficient) between LFP and VSD for all tetrodes and ROIs for the three periods. (C) (Left) Mean similarity across 20 trials between LFP and VSD for the three periods for all tetrodes and all ROIs (error bars represent SEM). (C) (Right) Mean similarity across trials between LFP and VSD for the same ROIs and tetrode for the three periods. (D) Spatiotemporal pattern of the mean evoked response (20 trials) to hind paw electrical stimulation. (E) Spike-triggered average (STA) maps are generated as the average of the VSD frames corresponding to the times when the corresponding neuron fired. (F) (Left) Example of STA VSD maps for one neuron from ROI 7 and one neuron from ROI 5 during baseline and evoked response periods. (Right) Average comparison between STA maps from same neurons and the average evoked response in (D) for baseline and evoked activity periods.

To measure the similarity between LFP and VSD signals we calculated the correlation coefficient for three periods: baseline (250 ms before stimulus onset), early evoked response (first 250 ms after stimulus onset) and late evoked response (250 ms after the early response) (Figure 4A). We calculated a similarity matrix between LFP signal from all the tetrodes and the VSD signal from all the ROIs for the three periods (Figure 4B). Note that the mean similarity between LFP and VSD signals increases significantly (t-test, p < 0.001) during the first 250 ms after stimulus onset, compared to baseline for all tetrodes and ROIs (Figure 4C left) and amongst same tetrode and same ROI (paired t-test, p < 0.05) (Figure 4C right).

Another study that is possible with our setup is to evaluate the involvement of single units in neuronal population activity after sensory stimulation. Evoked responses in primary sensory areas consist of two components. The first component occurs within the first 100 ms (early) after stimulus onset and the second component occurs during 150-400 ms after stimulus onset (late) (Sachidhanandam et al. 2013; Funayama et al. 2015; Manita et al. 2015). Consistent with this, we found that the evoked response in VSD, LFP signals and SUA were formed by these two components (Figure 3D-F and 4A). At the single-unit level, we observe that only neurons that were recorded close to the contralateral and ipsilateral HL cortical areas (neurons 4, 14 and 25 which come from ROIs 1, 5 and 8 respectively) fire within the early phase of the cortical response. However, most of the neurons recorded in the majority of remaining tetrodes participate in the late response (250-500ms after stimulus onset) (Figure 3D). In order to evaluate the participation of a single neuron in the functional ensemble, we calculated the spike-triggered average of VSD activity or STA maps (Figure 4E). We observe that when selecting spikes occurring during the baseline period of the stimulation trials, the STA maps do not show prominent activity compared to when the spikes are after the stimulus onset (evoked) (Figure 4F left). Unsurprisingly, the STA maps of neurons close to the somatosensory area (i.e. ROI 5), resemble the VSD evoke response closely (compare STA map of Figure 4F and evoked response in Figure 4D). However, even for some neurons distant from the hindlimb area (i.e. ROI 7. e.g. approx. 3 mm medial and 3 mm anterior to the center of the HL cortex), its STA map during evoked periods is significantly (t-test, p <0.01) more similar to the HL evoked pattern (Figure 4F right) than during the baseline period.

### Combined VSD imaging and multi-site electrophysiological recording of spontaneous cortical activity

The brain is constantly active, even in the absence of sensory input or motor output (Werner and Mountcastle 1963). We compared the LFP and the VSD signal when there was no stimulation. In general, VSD and LFP signals show similar dynamics as is observed in Figure 5A. However, this similarity varies with time and cortical location (Figure 5A). This type of variability has previously reported for small regions of visual areas during spontaneous activity (Arieli et al. 1995). However, with our setup, it is possible to extend this to wider regions of the brain. We measured the similarity between LFP and VSD signal for the same locations (tetrode and ROI) for 1 sec periods for 15 min of spontaneous activity. We observed that both signals have different levels of similarity depending on the recording location (Figure 5C). In addition, to compare the LFP and VSD signals, with our setup is possible to evaluate the participation of individual units from distant regions in cortical ensembles (STA maps). Previously, the relationship between VSD (or wide field imaging signals) and SUA has been studied by monitoring neurons from small regions of the brain (Arieli et al. 1995; Xiao et al. 2017). With our setup it is possible to study such relationship over large and distant areas. We observed that in most cases the STA maps are similar for the neurons recorded in the same tetrode (Figure 5B top row), however there are cases in which one can see different maps in the same tetrode (Figure 5B bottom row). This suggests that there are neighbouring neurons potentially participating in different ‘cortical ensembles’. Finally, we compare two STA maps for different neurons (recorded in different tetrodes) before and after each neuron fired (Figure 5D). We observe that in the top row case, in average, the neuron fires when a large population activity increases (time 0). In contrast, in the bottom row, the neuron fires after the large population activity starts to increase (time -20ms). This demonstrates that there is potentially a different pattern of cortical activity that might be related to different spiking patterns of different neurons (Sato et al. 2012).

**Figure 5.**
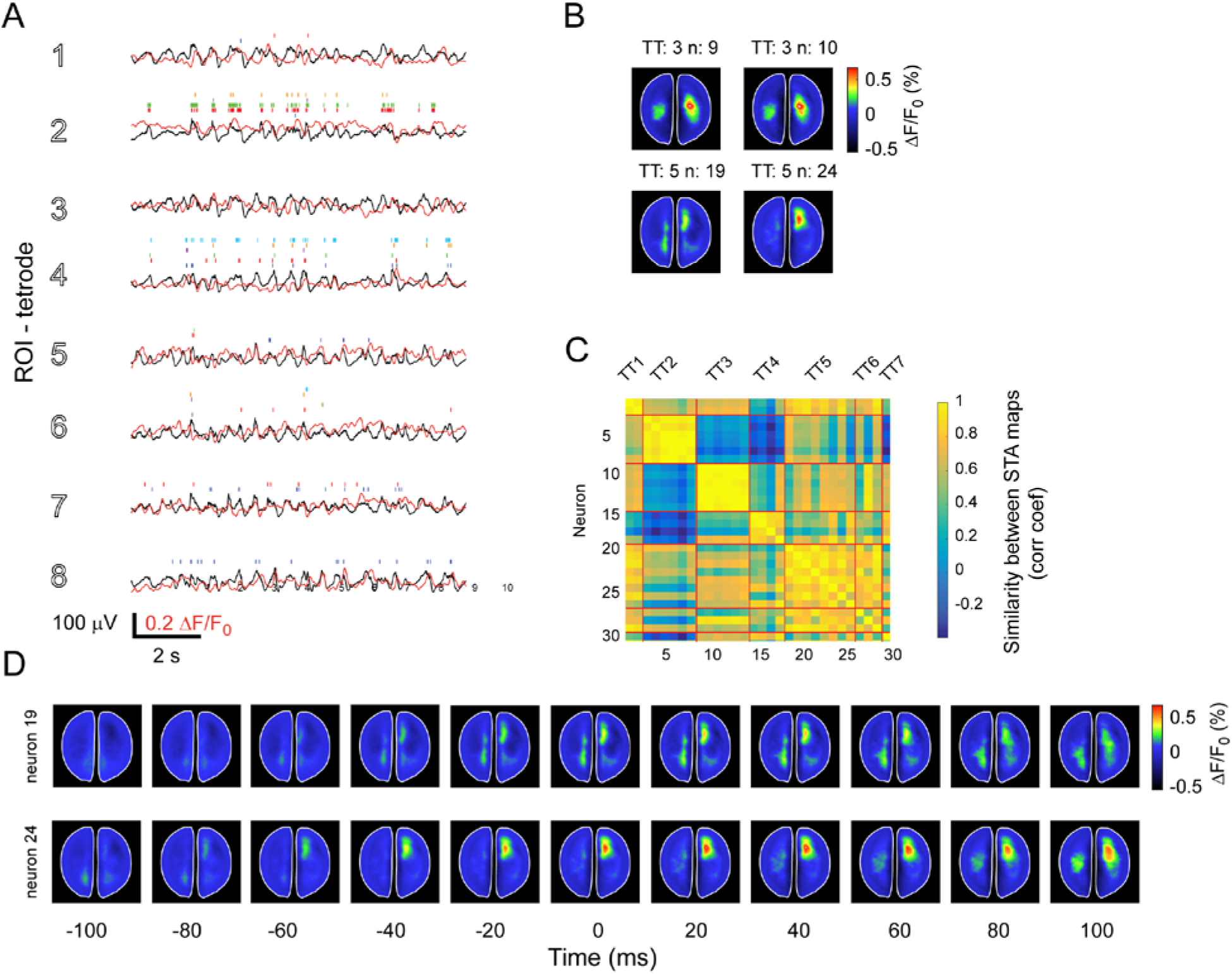
Simultaneous electrophysiological and VSD recordings during spontaneous activity in both hemispheres. (A) Simultaneous LFP, VSD and single unit recordings for the corresponding ROIs during 10 sec of spontaneous activity. The colors of the raster plots represent different neurons recorded in the corresponding tetrode. (B) STA maps of two neurons recorded in the same tetrode (TT) which are very similar (top row) and STA maps of two neurons recorded in the same tetrode which are different (bottom row). (C) Similarity matrix (correlation coefficient) for the STA maps generated for all neurons. (D) Example of STA maps for two different neurons (from different tetrodes) during spontaneous activity. The maps show the average activity 100ms before and after the neurons fired (time 0).

## 4. Discussion

Electrode arrays of different configuration have widely and successfully been used in neuroscience studies. Although these studies have greatly improved our understanding of cortical dynamics, their conclusions are limited due to the poor spatial coverage that causes a difficulty in monitoring neuronal activity across large cortical areas. In addition, the intrinsic limitation of electrophysiological signals in source localization due to volume conductance makes difficult to study contributions of particular cell types or brain regions (Katzner et al. 2009; Kajikawa and Schroeder 2011; Einevoll et al. 2013). Conversely, recent advances in protein based activity indicators such as voltage (Akemann et al. 2012; Gong et al. 2015; Abdelfattah et al. 2016), calcium (Chen et al. 2013; Bethge et al. 2017), glutamate (Marvin et al. 2013) sensors make possible to target specific type of neurons or locations in the brain. Such development in brain imaging technology has made brain imaging an extremely useful tool to monitor neuronal populations at a mesoscale level. Therefore, the combination of the multiple-site electrophysiology and VSDI represents a great opportunity to study brain function at different scales simultaneously. However, recording large neuronal population activity using a large amount of electrodes is rarely compatible with optical imaging due to technical difficulties. On the one hand, the requisite to provide independent movement and wire routing for each electrode or tetrode inevitably makes an electrode array bulky. On the other hand, wide-field imaging requires a large cranial window that needs to leave a clear space for excitation and imaging. As a way to resolve these issues, we developed a novel type of electrode array to combine deep cortical recordings with wide-field optical imaging (Figures 3 and 4). The array provides multisite electrophysiological recordings with arbitrary depth and a choice of electrodes to be used without interfering with wide-field imaging.

In this paper, we demonstrate the advantage of using our setup by simultaneously recording VSDI and electrophysiological data. With this method, it is possible to analyze the relationship between electrophysiological signals and VSDI recordings at different brain locations. In particular, we showed that, on average, the temporal dynamics between LFP and VSDI evoked responses are similar even for distant regions in the cortex (Figure 4). Analogously, we showed that during spontaneous activity the LFP and VSD signals are similar during 1 sec periods (Figure 5). Moreover, we showed that neurons that were recorded close to the HL S1 area (tetrode 5) participate in activity patterns that resemble the average HL evoked pattern (top row STA map in Figure 4F). However, even for neurons recorded far from the HL S1 area (tetrode 7), participate in population activity that vaguely resembles the HL evoked activity pattern (bottom row STA map in Figure 4F). This result highlights the advantage of combining multiple-site tetrode recordings and VSDI. Similarly, we demonstrated that, even though in most of the cases neurons recorded in the same location participate in a similar functional ensemble, there are cases where neighbour neurons (recorded in the same tetrode) are involved in different cortical networks (Figure 5B). Finally, we showed that the spatiotemporal dynamics of cortical activation patterns can be distinct even for neurons recorded in the tetrode (Figure 5D). This result demonstrates that spiking activity at single-cell level can be related to different neuronal ensembles at a population level, which is reflected in the STA maps (Sato et al. 2012; Okun et al. 2015). Further analysis using this approach could potentially clarify the relationship between cell assemblies (i.e. sequential activation of distinct neurons) at a local population level and cortical processing at a more global scale using wide-field imaging. Such relationships play an important role in top-down sensory processing (Roland et al. 2006; Zhang et al. 2014; Manita et al. 2015), in memory, in which sharp-wave events at a local hippocampal circuit might be related to more global cortical activity (Battaglia et al. 2004) or in cortical processing in general, as a mean to quantify neuronal ‘population coupling’ or ‘packet-based communication’ (Luczak et al. 2015; Okun et al. 2015).

In our proposed method to combine VSDI and electrophysiology there are important issues to note. We found that brain damage due to electrode loading inevitably leads to bleeding, which can compromise the quality of VSD imaging. Having *dura mater* removed permits easy rinsing of brain surface, whereas in dura intact preparations the extravasations may localize in subdural space. Among the electrodes we used in our experiments, we found the conventional tetrodes to be the most user-friendly. They are inexpensive and easy to make, and also hard to break. Conversely, sharp metal electrodes (Reitboeck 1983) have the advantage of producing less damage to the tissue, but they are considerably more fragile than conventional tetrodes.

In summary, we present a method to combine electrophysiological and wide-field imaging simultaneously. Such combination of techniques allows to monitor brain activity at different scales with high temporal resolution over large brain areas and it represents a great tool to study brain function in general (Ermentrout and Kleinfeld 2001; Grinvald and Hildesheim 2004; Takagaki et al. 2011; Mohajerani et al. 2013; Keane and Gong 2015). Therefore, the methodology presented in this paper further expands the available tools to improve the current understanding of brain function.

## Acknowledgements

This work was supported by Natural Sciences and Engineering Research Council of Canada (NSERC) Discovery Grant #40352 and #RGPIN-2017-03857 To MHM and BLM respectively, Campus Alberta for Innovation Program Chair (BLM & MHM), Alberta Alzheimer Research Program (MHM), Alzheimer Society of Canada (MHM). We thank Valery Bouquet for PCB design, Di Shao and Behroo Mirza Agha for animal breeding.

